# Blood donor biobank pipeline to collect genome-based samples for research

**DOI:** 10.1101/2025.09.30.679473

**Authors:** Jarno Honkanen, Veera A. Timonen, Jessica R. Koski, Julianna Juvila, Mikko Arvas, Blood Service Biobank, Finn Gen, Linnea Hartwall, Outi Kilpivaara, Rodosthenis S. Rodosthenous, Ulla Wartiovaara-Kautto, Arja Vuorela, Mark Daly, Aarno Palotie, Esa Pitkänen, Jukka Partanen

## Abstract

The integration of genome data with electronic health records, driven by large biobank studies, has advanced human genetics by allowing systematic exploration of genotype–phenotype links. Regular donation enables large, longitudinal sample cohorts. Because blood donors are generally healthy, disease treatments or progression do not disturb interpretations in functional studies. We describe here a pipeline on how to collect blood donors’ high quality plasma, serum, and living cell samples for multi-omics studies. Peripheral blood mononuclear cells (PBMC) were frozen and, after thawing, contained standard levels of immune cell subpopulations, responded to immune activation, and were of good quality starting material for multi-omics and cell imaging studies. We demonstrate that most genetic variants of interest to the major genomics study in Finland, FinnGen, could be found by random collection of samples during the standard blood donation without recall. Probing simple associations in the multi-omics data confirmed expected associations with e.g. age and sex, demonstrating good sample quality. As an example of interesting findings, we observed a significant association between frequent blood donation and lower levels of per- and polyfluoroalkyl substances (PFAS). The study demonstrates that regular blood donors are a suitable target population for high-quality, cost-effective sample collections.

## Introduction

Large biobank genome data collections combined with electronic health records have made phenome-wide association studies (PheWAS) feasible leading to increased power and novel discoveries in disease genetics^1–3^. Blood donors have been suggested to be an excellent option for large cohorts of healthy individuals as they voluntarily and repeatedly donate blood^4^. Based on in-depth interviews^5,6^, blood donors are known to have a positive attitude towards scientific research and use of their donated samples for research if not needed for patient care.

An example of a blood donor-based biobank^7^ is the Blood Service Biobank, organized by the Finnish Red Cross Blood Service. By the year 2024, more than 60,000 blood donors, encompassing nearly half of the blood donor population in Finland, have given the biobank consent that enables research use of biological samples and health registry data. The Finnish population has proved to be advantageous for genetic studies as it has encountered population bottlenecks, resulting in a relatively low genetic diversity and enrichment of some deleterious variants, as well as disease-predisposing and protective alleles^3,8^.

Following large-scale screening studies and identification of the disease-associated genetic variants, more detailed molecular and cellular studies are warranted for understanding their molecular effects and finding novel targets for drug development. For example, the FinnGen research project^3^ based on screening genome and registry data from 500,000 Finns has identified over a hundred novel genetic variants predisposing to or protecting from common diseases. Most of these variants are rare or even missing from other populations, but are found in Finland and sometimes in neighboring countries. Studies on the molecular effects of these variants could be done from samples collected by recalling patients or other individuals carrying the variants. Although samples of the patients with the disease linked to the variant certainly provide important advantages, such as disease-specific effects, they may be hard to recruit, and their variable disease progression and treatments may lead to heterogeneity or even masking of the exact molecular pathways affected. Regular blood donors provide an alternative or complementary source: they voluntarily and regularly donate blood, and as several serious illnesses such cardiovascular conditions and cancer restrict blood donation, they are absent of these diseases. Regular donation enables longitudinal sample collections for addressing variation within single individuals. We here describe a blood sample collection pipeline for obtaining good-quality peripheral blood samples for multi-omics molecular studies. As an example of the results obtainable from combining data from multiple sources, we found that higher donation activity was associated with lower levels of PFAS compounds.

## Materials and Methods

The overall aim of the FinnGen project^3^ is to improve our understanding of disease mechanisms and to enable the development of better treatment strategies for chronic, common diseases. To achieve this, FinnGen is analyzing correlations of genetic determinants enriched in the Finnish population and phenotype variation among the Finnish population. Whole genome analysis was performed for over 500,000 DNA samples from Finnish biobanks, and the genetic data were combined with the data obtainable from national health registries. GWAS analyses have revealed over 100 disease-associated gene variants^3^ that are strongly (the frequency at least twice higher in Finns as compared to non-Finns) enriched in the Finnish population. Understanding the molecular effects of these Finnish enriched disease-associated genetic variants was agreed to be of considerable interest to the FinnGen project. To this end, a sample collection pipeline called FinnGen 2 Expansion Area 5 (EA5) was set up to collect peripheral blood mononuclear cells (PBMC), plasma and serum biobank samples from 2500 healthy blood donors who have given a biobank consent and have earlier been genotyped in FinnGen.

Blood samples were collected from blood donors of the Finnish Red Cross Blood Service who had given a written biobank consent for the Blood Service Biobank of the Finnish Red Cross Blood Service, Finland. All samples were collected along with the standard blood donation; no specific recalls were done. The collection started March 8, 2021, and ended June 3, 2024. The aim was to process all samples within four hours from the time of collection at the donation site to ensure high quality. The overall pipeline is depicted in **Figure 1**, and details of the collection protocol are described in the **Supplementary Material**. Use of the samples and data is in accordance with the biobank consent and Finnish Biobank Act 688/2012. The FinnGen study has been approved by the Ethics Review Board of the Hospital District of Helsinki and Uusimaa (permit HUS/990/2017).

**Figure 1.**
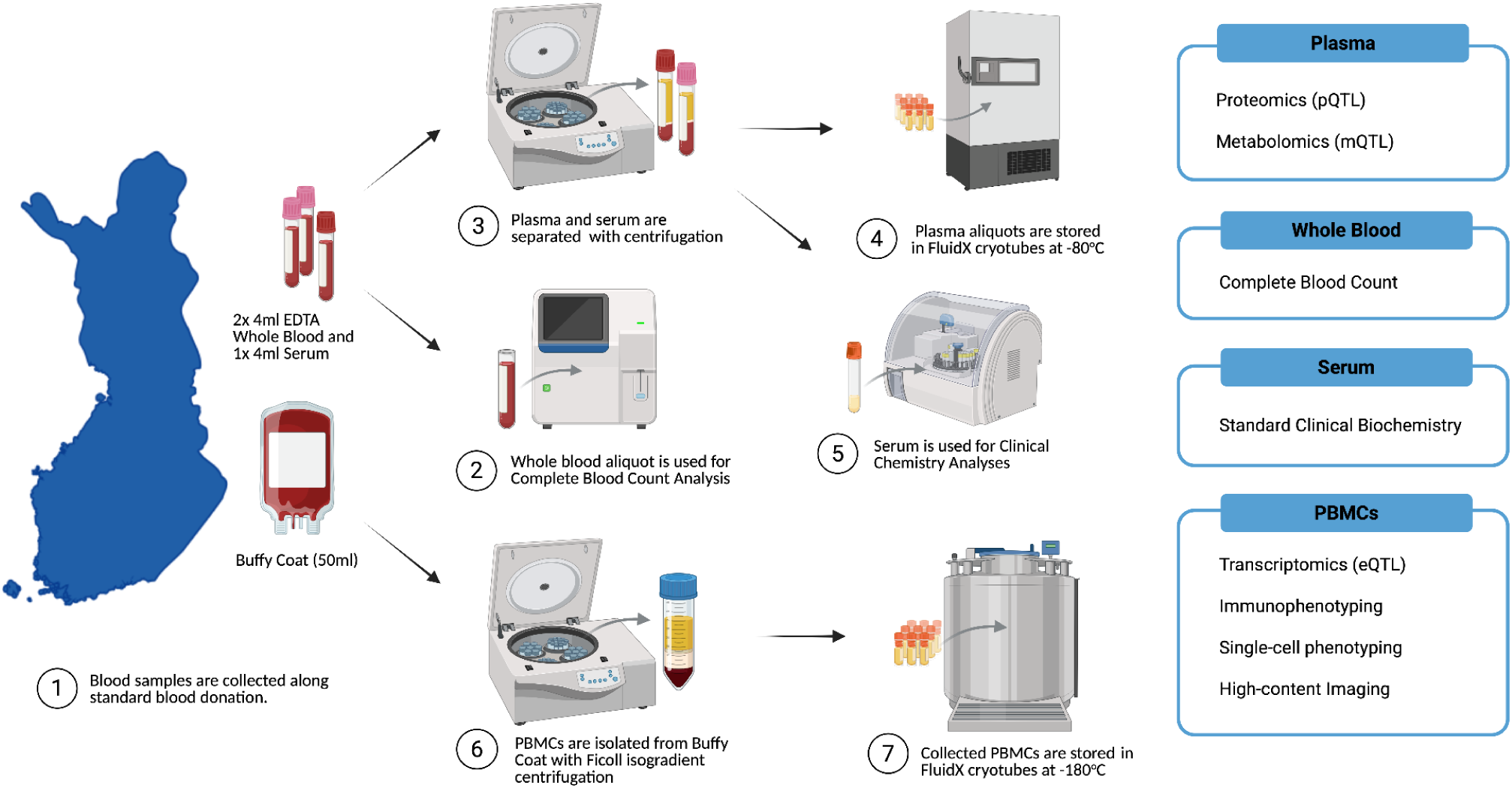
Overall pipeline of blood donor sample collection for biobanking, sample storage.

Details of the immune cell profiling and cytokine response, and analysis of the viability of frozen peripheral blood mononuclear cells (PBMCs) are described in the **Supplementary Material**. Briefly, the immune cell populations were classified based on staining using anti-CD4, anti-CD141, anti-CD123, anti-CD11c+, anti-HLA-DR, anti-CD127, and anti-CD25 antibodies. Analysis of the following six immune cell subpopulations was performed with BD Influx™ (BD Biosciences): Teff (CD3+CD4+CD25intCD127+), Treg (CD3+CD4+CD25hiCD127-), mDC1 (HLADR+CD11c+CD141-), mDC2 (HLADR+CD11c+CD141+), pDC (HLADR+CD123+CD4+), and monocytes (CD4lowHLA-DR+). Information on the fluorochrome-conjugated antibodies used for the FACS analysis of different immune cell populations is shown in **Supplementary Table S2**.

To demonstrate the functionality of the cell samples, the thawed PBMCs were stimulated for 48 h with plate-bound anti-CD3 (BD Pharmingen) and soluble anti-CD28 (1 µg/ml) (BD Pharmingen). Six replicates of 2 x 10^5^ PBMCs were performed. After the stimulation, the cell culture supernatant was collected for cytokine analysis. The concentrations of cytokines, chemokines, and growth factors were analyzed using the 38-plexed Milliplex MAP Kit (cat.no. HCYTMAG-60K-PX38) according to the manufacturer’s recommendations (Merck-Millipore Corp., Billerica, MA, USA). The kit measured the levels of the following molecules: sCD40L, EGF, Eotaxin, FGF2, FLT3L, Fractalkine, GCSF, GMCSF, GRO, IFNalpha, IFNgamma, IL1alpha, IL1beta, IL1RA, IL2, IL3, IL4, IL5, IL6, IL7, IL8, IL9, IL10, IL12p40, IL12p70, IL13, IL15, IL17A, IP10, MCP1, MCP3, CCL22, MIP1alpha, MIP1beta, TGFalpha, TNFalpha, TNFbeta, and VEGF.

## Full blood counts and clinical chemistry

As a part of the EA5 pilot, complete blood count (CBC) and clinical chemistry analyses were obtained from the plasma samples of blood donors (n = 2200 samples) (https://www.finngen.fi/en/other-biological-data). CBC includes measurements on the amounts and sizes of red blood cells, hemoglobin, platelets and white blood cells. Clinical chemistry analysis measures compounds in blood, such as cholesterol.

## Metabolomics

The samples were frozen within four hours after bleeding and stored at −80°C. Sample preparation was automated, including protein precipitation using methanol, followed by centrifugation. Extracts were divided into fractions for analysis under various chromatography and ionization methods: two reverse-phase UPLC-MS/MS with positive ion mode, one with negative ion mode, and one HILIC with negative ion mode. Quality assurance included spiked internal standards, pooled technical replicates (CMTRX), blanks, and solvent controls to assess and monitor variability across sample processing and instrumentation.

High-resolution mass spectrometry (UPLC-MS/MS) was conducted using a Thermo Q-Exactive Orbitrap system. Peak identification relied on a comprehensive library of over 3,300 authenticated standards and robust criteria, including retention index, accurate mass, and MS/MS spectral matching. Data processing involved stringent QC, curation to exclude artifacts, and quantification based on peak area. For multi-day analyses, batch effects were corrected using a block-correction method. Bioinformatics tools integrated LIMS tracking, automated data extraction, and visualization platforms to ensure high data fidelity for downstream statistical and biological interpretation.

## Proteomics

The aptamer-based proteomic platform SomaScan® 7K Assay v4.1 (SomaLogic, Boulder, CO, USA) was used to quantify approximately 7,000 proteins from a minimal sample volume of 55 µL of EDTA plasma. The assay utilizes SOMAmer® (Slow Off-rate Modified Aptamer) reagents, which are single-stranded DNA aptamers engineered for high specificity and affinity to their target proteins. Before assay, plasma samples underwent a series of processing steps, including dilution and incubation with SOMAmer reagents. Post-binding, unbound proteins and non-specifically bound reagents were removed through bead-based immobilization and washing steps. The bound SOMAmer-protein complexes were then isolated, and the SOMAmer reagents were quantified on microarrays.

Data normalization was performed through a multi-step process designed to eliminate systematic biases and ensure consistency across samples and assay plates, as described in SomaScan^®^ Assay v4.0 Technical note (https://somalogic.com/tech-notes/). Each sample’s data was scaled using a factor derived from the median ratio between control relative fluorescent units (RFU) and a reference within the same plate. This was followed by Intraplate Median Signal Normalization, applied only to calibrator samples. A scale factor for each SOMAmer was calculated by dividing the global reference RFU by the local calibrator RFU on each plate, and this factor was applied across the entire plate. Median Signal Normalization to a reference was performed on all individual, QC, and buffer samples by comparing each to a global reference dataset from healthy individuals. This step applied separate scale factors for each dilution group per sample, ensuring consistency across the full dataset. Together, these normalization steps reduced assay variability and standardized the data for robust downstream analysis.

Olink proteomics was done essentially as described by Argentieri et al^9^. The study included a set of samples from the cohort described in the present study. Briefly, the frozen samples were shipped to the Olink Bioscience Laboratory (Uppsala, Sweden) for analysis using the 3,072 multiplex proximity extension assay. The sample plates were normalized using an internal extension control and an inter-plate control. Negative control samples without antigen were included.

## Association analysis of metabolomic, proteomic and clinical data modalities

Association analyses were conducted for all studied metabolites (n = 722 donors; n = 1236 metabolites), proteins from Olink (n = 934 donors; n = 2836 proteins), blood tests (n = 1159 donors; n = 26 parameters), and clinical chemistry tests (n = 1092 donors; n = 16 parameters) available. All values used in analysis were concentrations or amounts of the studied molecules. The following covariates were used to form the linear model: *y* ∼ age in years + sex + ancestry principal components (PCs) + Body Mass Index (BMI) + smoking status (of ever having smoked) + year of collection + month of collection + hour of collection + batch covariates + blood donation count in the last two years + time between donation and centrifugation + time between centrifugation and freezing. Two of the latter time covariates were reported in minutes. Ancestry PCs were available from FinnGen and demonstrate the ancestral background of donors. Batch information was available for proteomics data, in which case the batch covariates included plate, row, column and batch, as the proteomics data was created in three separate batches. Month and hour were transformed to sine and cosine measurements as a circular representation of time using the following formulas.

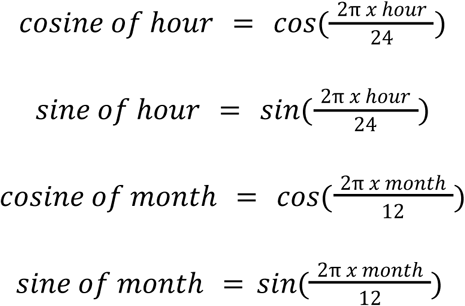

Proteomics and metabolite level values were log-transformed. For all data modalities (protein, metabolite, CBC and clinical chemistry measurements), values were scaled to a mean of zero and standard deviation of one. For covariates age, BMI, blood donation frequency, time between donation and centrifugation and time between centrifugation and freezing were scaled to a mean of zero and standard deviation of one to achieve comparability for analysis. For each analyte *y,* a least-squares linear regression model was fitted. Benjamini-Hochberg procedure was applied to the resulting *p*-values to adjust for multiple testing, separately for each data type of proteomics, metabolomics, blood tests and clinical chemistry tests. Analyses were performed in R in the FinnGen Sandbox environment using base R^10^ (version 4.5.1), and visualizations generated with packages such as ggplot2^11^, and seaborn^12^ in Python.

## Bayesian analysis of metabolomic data for blood donation frequency

To better estimate effect sizes, Bayesian association analysis was employed for a subset of metabolites that gave a significant association result of false discovery rate (Benjamini-Hochberg) < 0.01 for blood donation frequency. The covariates were the same as with the initial linear modeling (**Methods**), with the addition of interaction between sex and count of donations in the last two years (*y* ∼ age + sex + ancestry principal components (PCs) + BMI + smoking status (of ever having smoked) + year of collection + month of collection + hour of collection + batch covariates + blood donation count in the last two years + time between donation and centrifugation + time between centrifugation and freezing + sex*blood donation count in the last two years). Bayesian analyses were performed with R using the package rstan^13^.

## Results

To investigate the functional effects of genetic variants that are both enriched in the Finnish population and associated with various multifactorial diseases, we established a pipeline to collect fresh blood samples (plasma, serum, and PBMCs) from healthy donors during routine blood donation. Samples were obtained exclusively from donors who had provided biobank consent and whose genomic data had been analyzed in the FinnGen project^3^ and subsequently returned to the Blood Service Biobank (hereafter referred to as the Biobank). For logistical reasons and to ensure rapid processing and freezing of samples post-donation, recruitment and sample collection were primarily done in the Helsinki metropolitan area. Due to internal migration trends toward the capital area of Finland^14^, this region was expected to represent the general Finnish population. Additionally, smaller-scale collection sites were piloted at blood donation centers in two other locations: Oulu in northwestern Finland and Kuopio in the northeast.

The pool of potential blood donors fulfilling the above-mentioned inclusion criteria in the entire Finland was ∼55,000, of whom ∼8,500 had donated during the year 2023 in at least one of the three donation sites in the Helsinki capital area. The target for resampling was set at the present pilot to 2,500 (FinnGen Extension Area 5 or EA5 project). The samples (**Fig. 1**) were taken during the standard whole blood donation from the diversion pouch without active recalling. The buffy coat fraction was saved for PBMC preparation. It is of note that the production of the standard red cell and platelet units could be done from these donations. The overall pipeline and sample collection are depicted in **Figure 1**. The samples were aliquoted and frozen for downstream analyses, such as multi-omics studies of the variants.

Amongst the first 2,500 donors of whom a re-sample was obtained, 98% of the 136 variants that were disease-associated and Finnish-enriched^3^ were found at least once in a heterozygous state (**Fig. 2A**) and 40% of the 136 variants could be found at least once in a homozygous state (**Fig. 2B**). For 88% of the variants more than five heterozygous or homozygous carriers were found. For the variants with an allele frequency of below 1%, a targeted approach may be needed.

**Figure 2.**
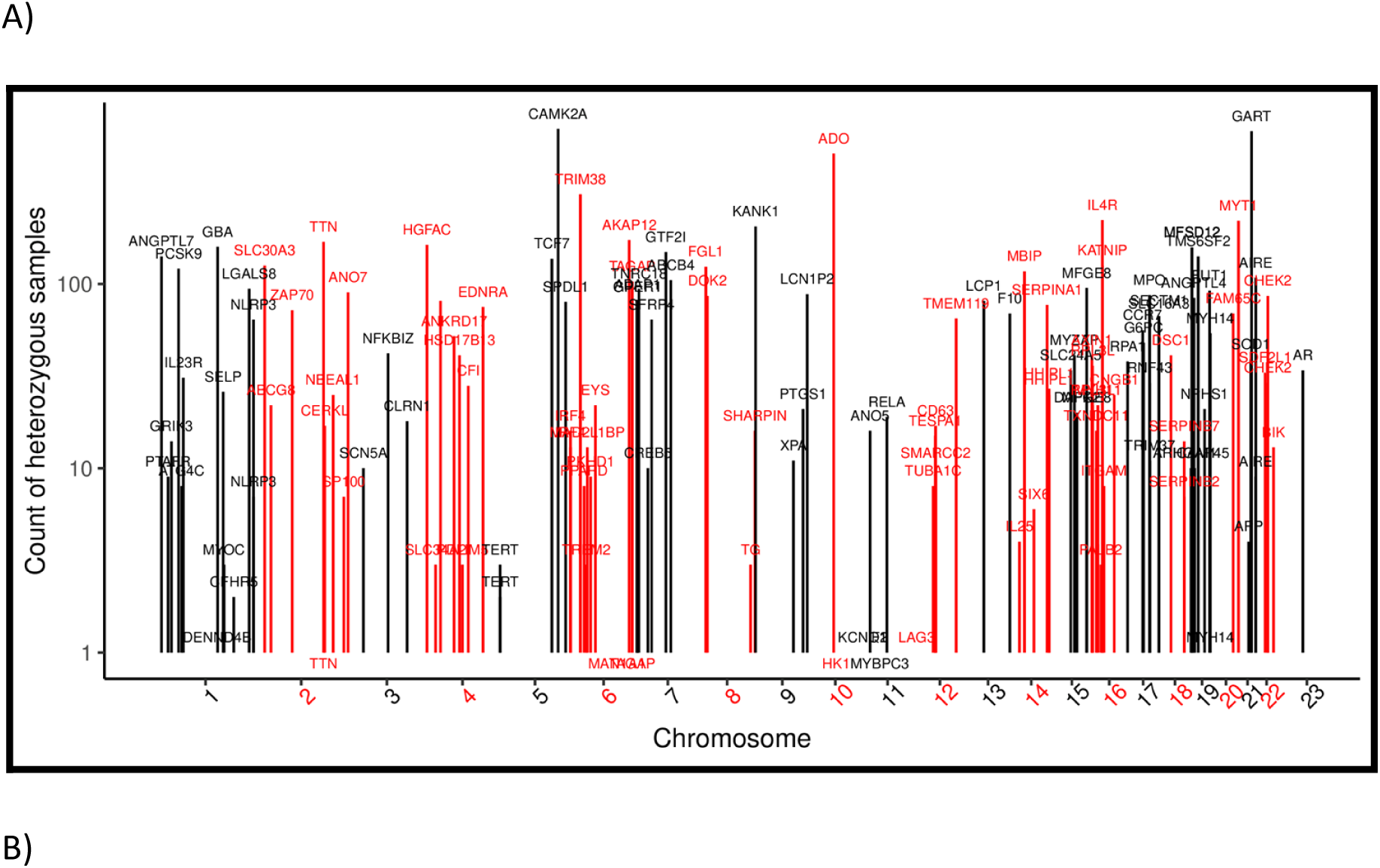
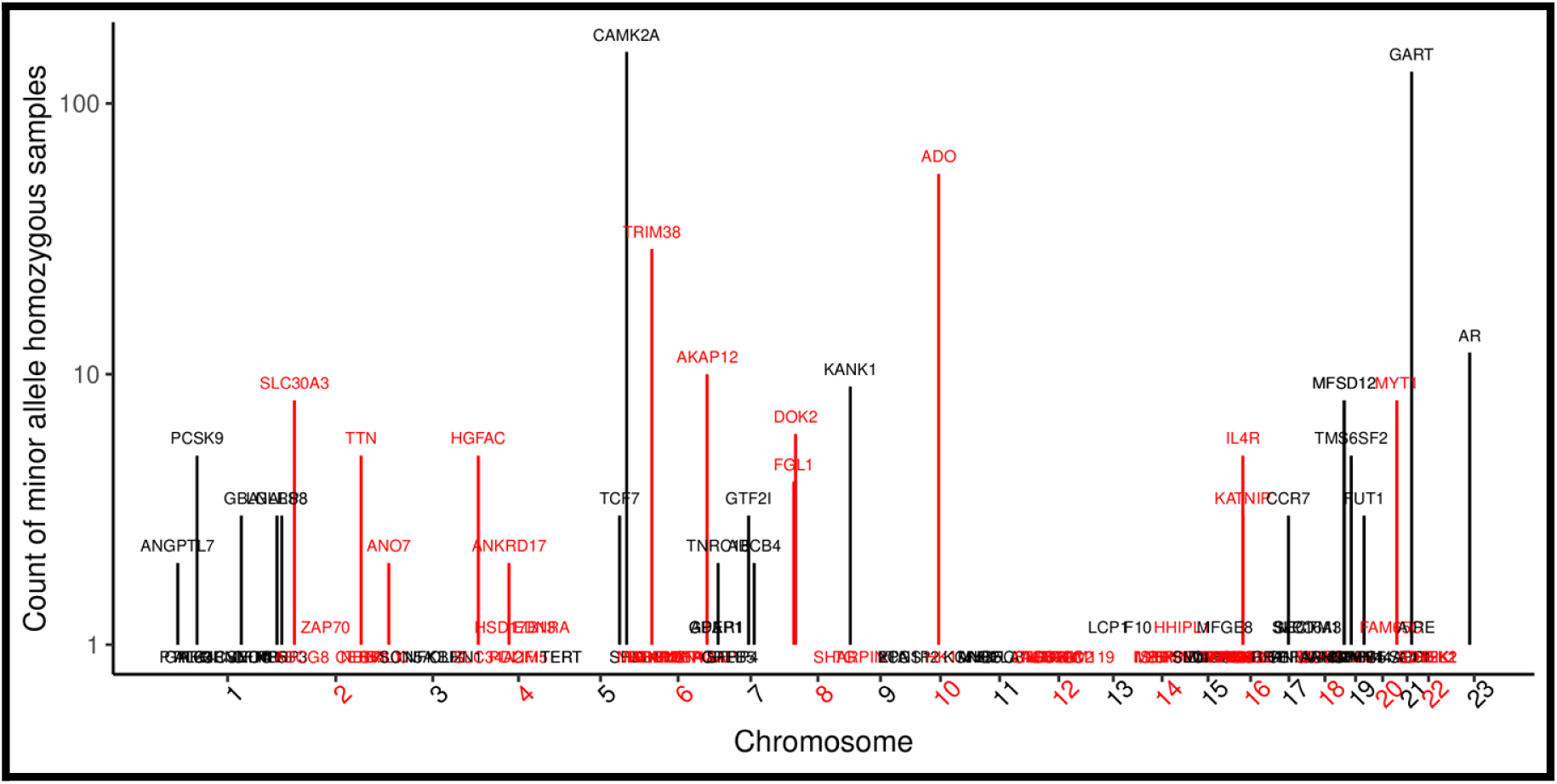
A) The number of heterozygous carriers of 136 target variants found to be associated with disease in the FinnGen project among the 2,500 blood donors. B) The number of homozygotes.

High sample quality is essential for downstream analyses of biobank samples, particularly for complex and expensive multi-omics studies. **Figure 3** shows an example of immune cell populations obtained from the thawed PBMC samples using antibody staining. In the four samples tested, the levels of the major immune cell populations obtained were as expected (**Suppl. Fig. 1**). The PBMCs showed a good response to the anti-CD3 + anti-CD28 antibody activation and produced the expected cytokine profiles. The concentrations of key T cell cytokines in untreated and stimulated PBMCs are shown in **Figure 3B**. The concentrations of other cytokines are shown in the **Supplementary Table 3**. Metabolomics analysis is known to be sensitive to the sample quality. One thousand (n=1000) plasma samples were analyzed with the Metabolon global discovery panel. Overall, 1,602 metabolites were reported by the manufacturer to be detectable, with over 1,200 of these known, named metabolites. In the Olink and SomaScan® proteomics platforms, the samples also showed a high quality, with over 2,900 proteins reported to be detectable.

**Figure 3.**
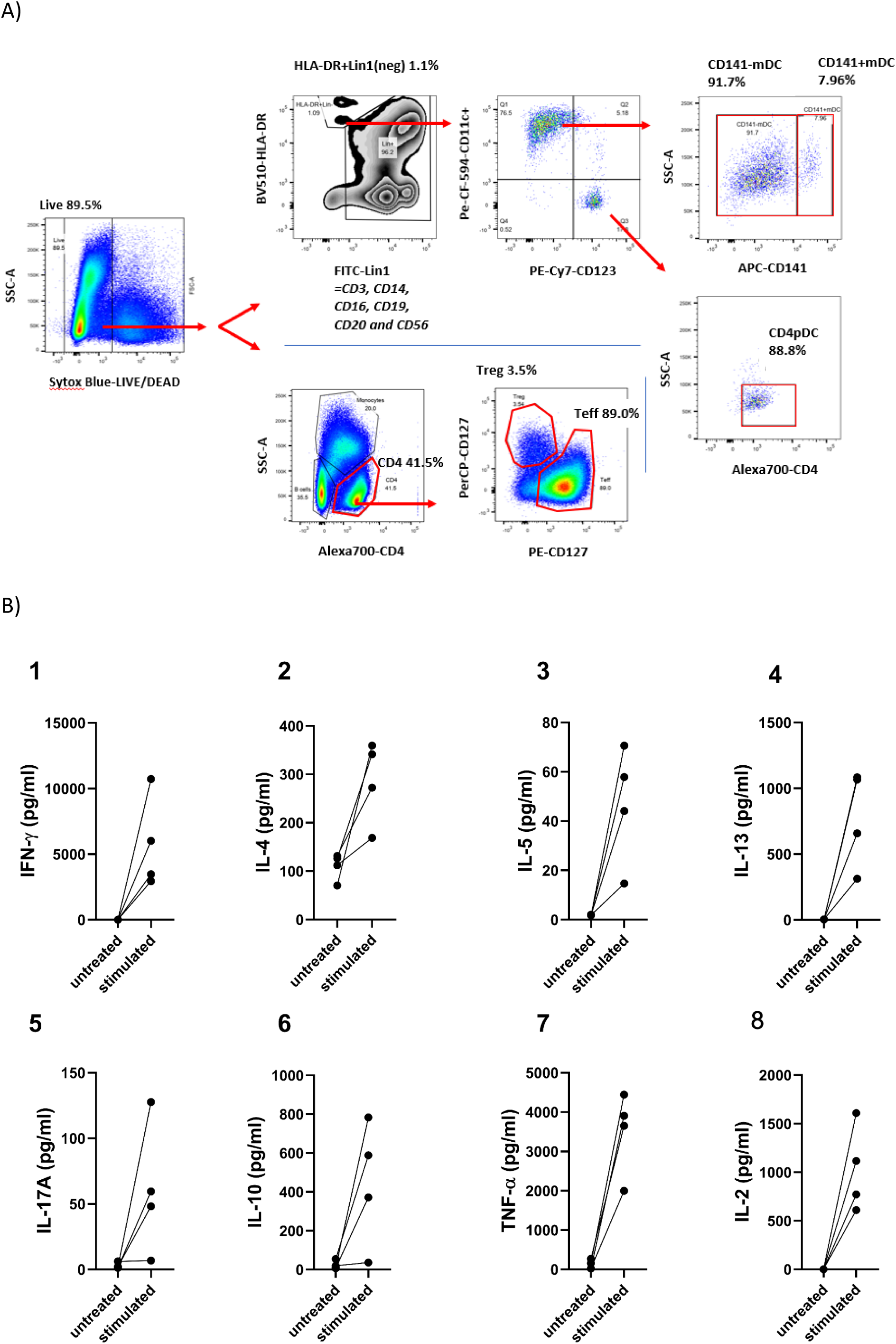
A) Identification of effector T helper cells (Teff), regulatory T helper cells, CD141(-)mDCs, CD141(+)mDCs, pDCs, and monocytes in the PBMC samples thawed after freezing. The dead cells were excluded from the analysis by the Sytox blue staining. Viable cells were gated based on the cell surface antigen expression. B) Production of key T cell cytokines by thawed PBMCs after anti-CD3 + anti-CD28 stimulation. Cytokine levels (pg/ml) in supernatants are shown for individual donors under untreated and stimulated conditions. Each plot displays concentrations of a specific cytokine: (1) Th1 associated IFN-γ, (2-4) Th2 associated IL-4, IL-5 and IL-13, (5) Th17 associated IL-17A, (6) regulatory T cell cytokine IL-10, (7) cytotoxic T cell associated TNF-α and (8) cytokine promoting growth, differentiation and survival of T cells, IL-2. Each dot represents one donor, with lines connecting paired responses from the same individual.

Figure 4A shows the data modalities completed from the study cohort at the time of writing the present paper (year 2025). Gender, age, and other cohort demographic data can be seen in Figures 4B-E. To understand whether the sample quality is sufficient to carry out high-quality research, we probed associations in the multiomics data available in FinnGen (Fig. 5) between measurements including blood counts (26 parameters), clinical chemistry (16 parameters), proteomes (>2,800 proteins) and metabolomes (>1,200 metabolites), and donor and sample parameters including demographics, blood donation counts and sample processing times (donation–centrifugation–freezing time). Significant associations were seen particularly with sex, age, smoking, and body mass index (BMI) (**Suppl. Tables S4-S9**).

**Figure 4.**
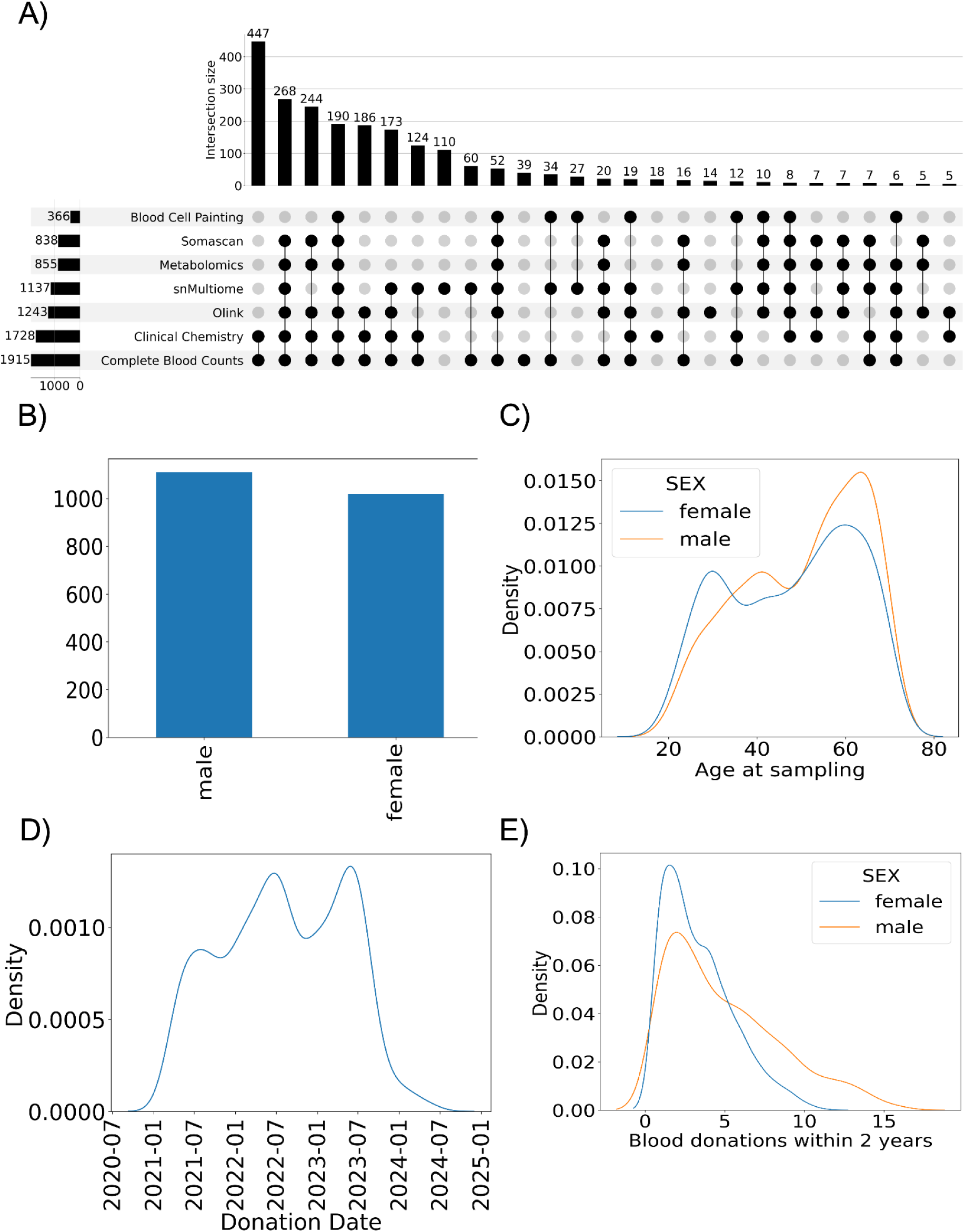
Overview of the omics and donor characteristics in the EA5 dataset. A) The number of samples analyzed for each of the seven omics data types and their combinations. Combinations with less than five donors are omitted to conform with FinnGen’s data policy. Basic blood donation records and demographics were available from all 2,500 blood donors. B) Sex distribution. Density plots for C) age at sampling, D) sample donation date, and E) number of blood donations within two years of sample donation.

**Figure 5.**
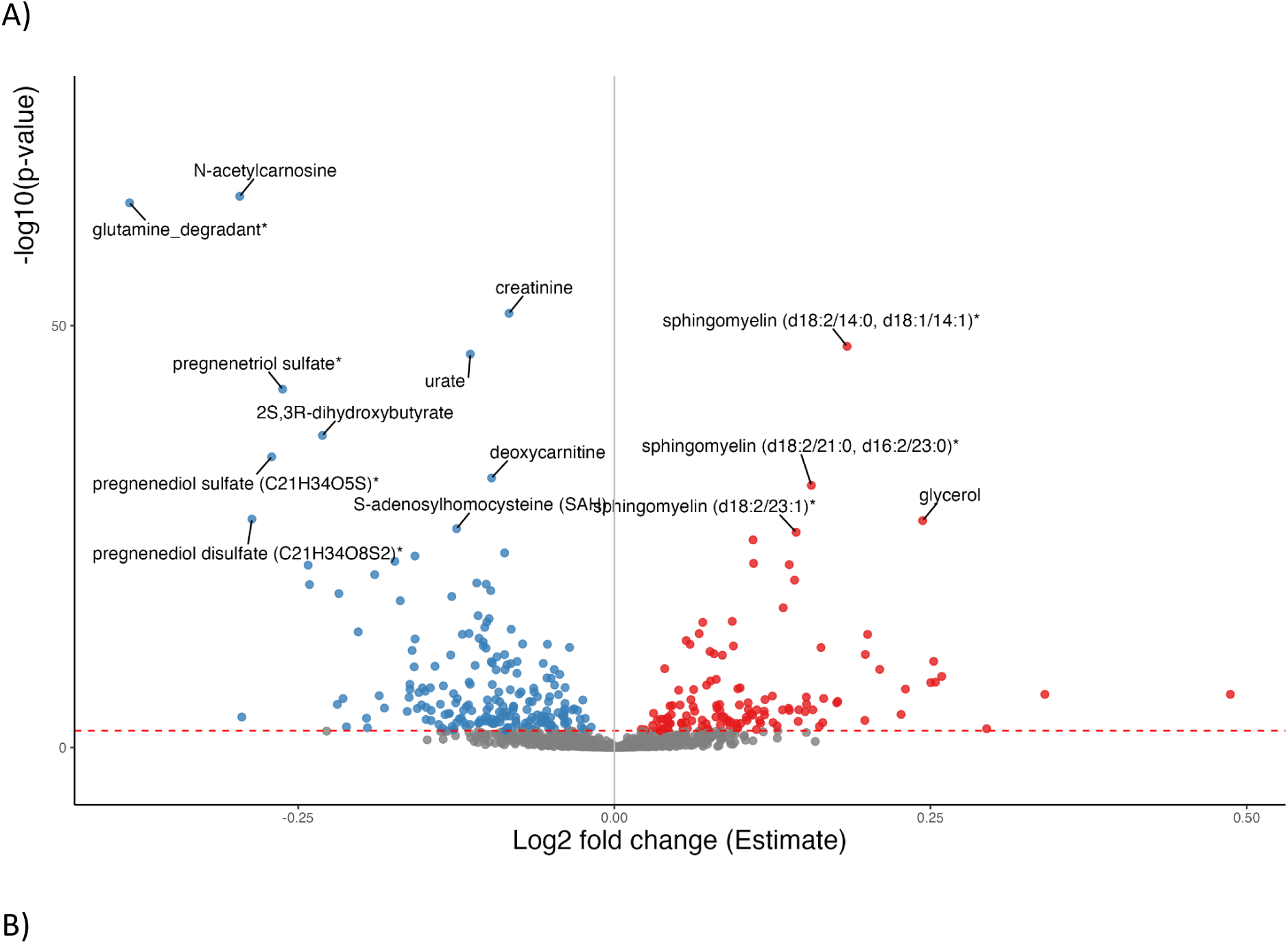
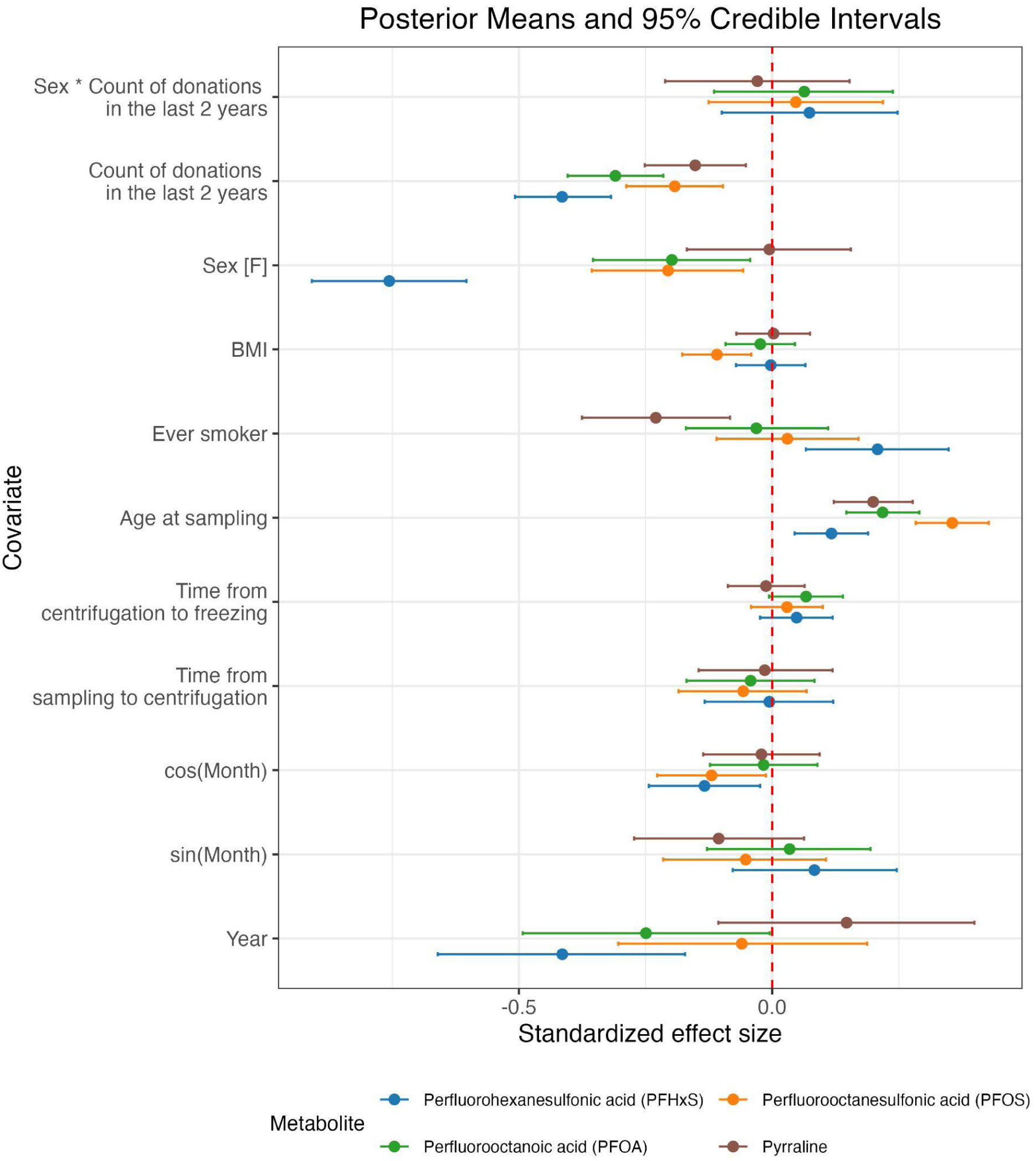
A) A volcano plot of associations between sex of the donors and metabolite levels (n=722 donors, FDR=0.01 shown as a red line). Log2 fold change (Estimate) on the x-axis indicates the direction of the effect: positive estimates indicate higher values of metabolite levels for female donors compared to male donors. B) Bayesian association analysis for the levels of PFAS compounds (PFHxS, PFOS, PFOA) and pyrraline, as well as donor and sample characteristics. Compound levels were associated negatively with blood donation frequency, and positively with age at sampling. Female sex was negatively correlated with PFHxS in particular.

Levels of many metabolites were associated with sex (**Suppl. Fig. 2 A-C**), age, and BMI, compatible with previous findings15,16. For instance, 5α-Androstan-3α,17*β*-diol 17-glucuronide, an androgen-derived metabolite more abundant in males, showed a strong association with sex (*β*=-1.36; FDR=8.3×10^−94^). Smoking status was also associated with *e.g.* cotinine (*β*=0.59; FDR=9.17x10^-12^), a metabolite of nicotine, while glutamate levels were positively associated with BMI (*β*=0.37; FDR=2.1x10^-24^). Blood donation activity was found to be associated with PFAS levels in both linear regression and Bayesian analyses (**Suppl. Table S4**, Fig. 5B, **Suppl. Fig. 3**), showing that more whole blood donations over the past two years were associated with lower PFAS concentrations, whereas older age was associated with higher levels. The clearest sex-specific difference emerged for PFHxS, where the blood donation frequency correlated with lower reduced PFHxS levels in both male (*β*=−0.41, FDR=3.78x10^−9^) and female donors (*β*=−0.35, FDR=6.97x10^−5^). Whereas there was a significant correlation between PFHxS levels and age in female donors (*β*=0.27, FDR=5.64x10^−8^), no such correlation was observed in male donors (FDR=0.77), in contrast to earlier studies^17^ (**Supplementary Table S5, S6,** Figure 5B). However, the relationship between PFAS concentrations and age has been suggested to be non-linear between sexes^18^.

In plasma proteomics, similarly to metabolomics, levels of multiple proteins correlated with sex, age, and BMI (**Suppl. Fig. 2 D-F**). BMI and leptin (*β*=0.51; FDR=2.62x10^−89^) were correlated, as leptin has an established role as a hormone released from adipocytes controlling energy expenditure and appetite (**Supplementary Table S7**). For example, INSL3 (*β*=-1.5; FDR<2.225x10^-308^) and SPINT3 (*β*=-1.6; FDR<2.225x10^-308^) levels were increased in male donors. These proteins are known to be male-specific and expressed mostly in gonadal tissues^19^. Transferrin levels were associated with the number of blood donations (*β*=0.17, FDR=0.00046), as was expected due to iron loss in donations. It is of interest that the time between donation and centrifugation was associated with an increase in levels of multiple proteins. Some of these, such as cytokines IL1B (*β*=0.28, FDR=1.3x10^-12^) and TNFSF14 (*β*=0.19, FDR=3.2x10^-7^), as well as the CD84 membrane protein (*β*=0.17, FDR=1.5x10^-5^) (**Suppl. Table S7**) are related to the immune response. Additionally, multiple proteins were found to lower towards the evening (**Suppl. Table S7**; e.g. CD244 and cosine of hour *β*=-4.26, FDR=1.4x10^-5^). Cosine of hour represents the cyclical nature of time, and reaches its highest value at midnight (**Methods**).

## Discussion

The present study shows that the blood donor population who voluntarily and often regularly donate whole blood can provide an excellent source of biological samples for scientific research. There are many aspects supporting this conclusion. Importantly, in-depth interviews and sociological studies have shown that the attitudes of blood donors are very research-prone, and they readily accept leftover blood samples to be used for research4–6. The studies^5,6^ indicate that open information and mutual trust are fundamental for research use of blood donor samples. The national biobank law^20^ provides the legal framework for collecting samples and gives guidelines for consents, information, and regulation. As shown in the present study, research samples can readily be obtained during the standard blood donation; there is no need for research-only invitations for standard small-volume samples. Existing blood donation sites have been organized for safe and competent blood drawing, there is no need for each project to set up their own costly sample collection facilities. This results in a cost-effective way of collecting samples. The sample collection can readily be organized nationwide as blood services usually collect blood from multiple sites across the operating area or country. In addition, the blood drawn into the so-called diversion pouch - the blood sample taken for virus and blood type testing in each whole blood donation – contains all major cell populations and plasma; the volume drawn, albeit a limiting factor, usually suffices for many research purposes. Red cells, platelets, and plasma are the major products for transfusion medicine, while no living PBMCs are needed. Hence, PBMCs, even from the entire blood bag (∼500 ml), can be collected for research purposes without reducing blood supply for patients. We here confirm that the PBMC fraction indeed survives freezing and thawing. A large-scale alternative to a diversion pouch for PBMCs are leukocyte filters^21^ used for the leukoreduction of blood products from which living PBMCs can be eluted. An obvious limitation to blood donor samples is restriction to blood-derived samples only. Another caveat may be the healthy donor effect^22^: continuous selection of particularly healthy individuals as regular donors, most likely also leading to genetic bias.

The blood samples of the present study were primarily collected for the FinnGen research project^3^ for extensive multi-omics analyses to understand functional consequences of disease-associated gene variants. Blood donors are relatively healthy but can provide a promising sample source for mechanistic studies of disease-associated gene variants as there should not exist variation caused by variable disease progression and therapy protocols between patients. Naturally, samples from patients also are essential for many studies and can be collected from hospitals.

The present study demonstrated that we could detect the expected associations of metabolites and proteins with basic parameters, such as age, BMI, and sex. These results confirm that the sampling protocol resulted in samples of sufficient quality for extensive omics studies. There were a few findings that are of practical interest and that must be considered as possible confounding factors in sampling protocols. Firstly, the time between donation and centrifugation was associated with an increase in the levels of multiple proteins, which may result in artefacts when analysing samples with heterogeneous sampling protocols. Among these proteins were the cytokines IL1B and TNFSF14, and proteins related to immune regulation, such as CD84 and LRCH4. In addition, plasma levels of TEK receptor tyrosine kinase and SMNDC1 splicing factor were associated with this time interval. Multiple proteins were significantly affected by the time of day, which could indicate circadian variation affecting protein levels. However, all samples were collected during the workday, so the cyclical modeling will only show effects that occur during the day and may thus distort real effects.

The frequency of blood donation was associated with transferrin levels, which was expected as whole blood donation results in iron loss that needs to be compensated from iron stores. Metabolomics showed that frequent blood donation reduced the plasma levels of PFAS compounds. The PFAS compounds are known to be persistent and harmful substances. The finding that their plasma levels can be reduced by whole blood donation is expected and has been reported in firefighters^23^ who are known to have high PFAS levels. Gasiorowski *et al*.^23^ reported that blood and plasma donations resulted in lower PFAS levels than the control groups. In high-risk populations, drawing blood or plasma could be a way to reduce high PFAS loads. Olsen *et al*. showed a decline in PFAS levels in American blood donors in respect to time, but suggested it to be the result of reduced exposure in the areas where the samples were taken^24^. There are no specifications for PFAS levels in blood products and the long-term effects and levels of PFAS substances transferred from donor to recipients of blood products are not known. Transfer of PFAS substances, however, can be considered not to be a critical issue in transfusions needed in emergencies to save patients’ lives.

This cohort was collected into the Blood Service Biobank. According to the Finnish biobank act, these samples and data can be requested for research purposes from the Biobank. In addition to the blood samples described here some omics results and imputed complete blood types^25^, classical HLA^2^6, non-classical HLA^27^, and KIR genotypes^28^ of these samples are available from the biobank. High content image analysis with the cell painting assay has been described by Högel-Starck *et al*.^29^. The samples are analysed for extensive multiomics including joint single-cell gene expression and chromatin accessibility profiling (Kanai *et al. unpublished*) in the FinnGen research project and these results may be available for studies by contacting the FinnGen project^3^ (www.finngen.fi). In conclusion, we here demonstrate that high-quality blood donor samples for multiple research purposes can be readily collected along the standard blood donation. Biobank consent and the Finnish biobank legislation enable not only the distribution of samples and data to the research projects but also combining the analysis results with health registries.

## Acknowledgements

This study was supported by funding from the Blood Service Research Fund (to E.P.), Finnish Cancer Foundation, Sigrid Juselius Foundation and Academy of Finland (to O.K.), and FinnGen. The costs of collecting the 2500 samples were partially funded by the FinnGen research project. We want to thank all biobank blood donors and the staff of blood donation sites for their invaluable help, as well as Lea Urpa, Essi Viippola and other colleagues for discussions. BioRender.com was used in creating figures.

## Supplementary Figures

**Supplementary Figure 1.**
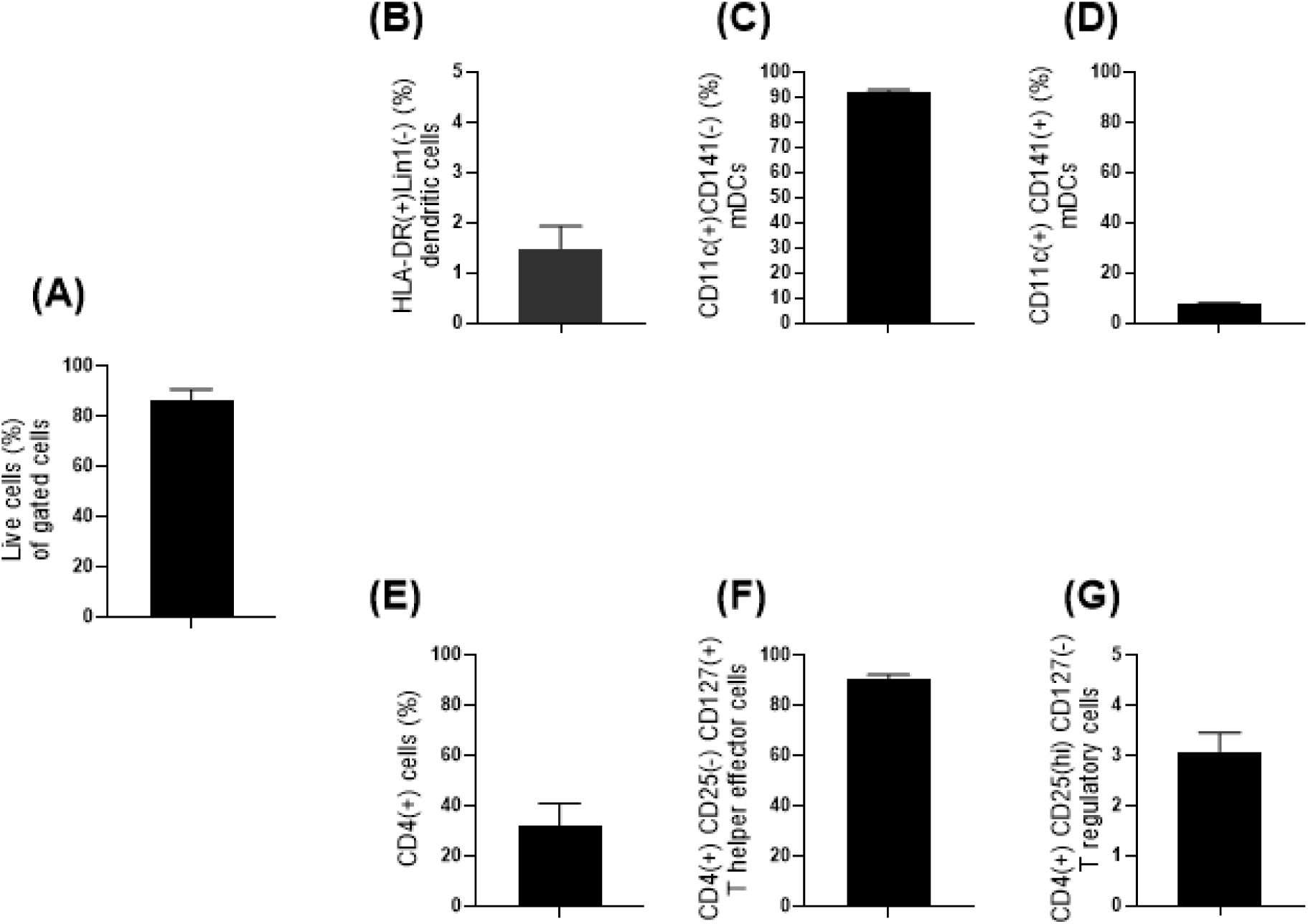
Analysis of cell viability and frequencies of dendritic cell and T helper cell subpopulations in thawed PBMCs from four blood donors. Panel A: On average, 86.1 % of gated cells were viable after thawing. Panels B–D: Frequencies of HLA-DR^+^ Lin^−^ cells (dendritic cells), CD141^−^ mDCs, and CD141^+^ mDCs. Panels E–G: Frequencies of CD4^+^ T helper cells, T helper effector cells, and T regulatory cells.

**Supplementary Figure 2.**
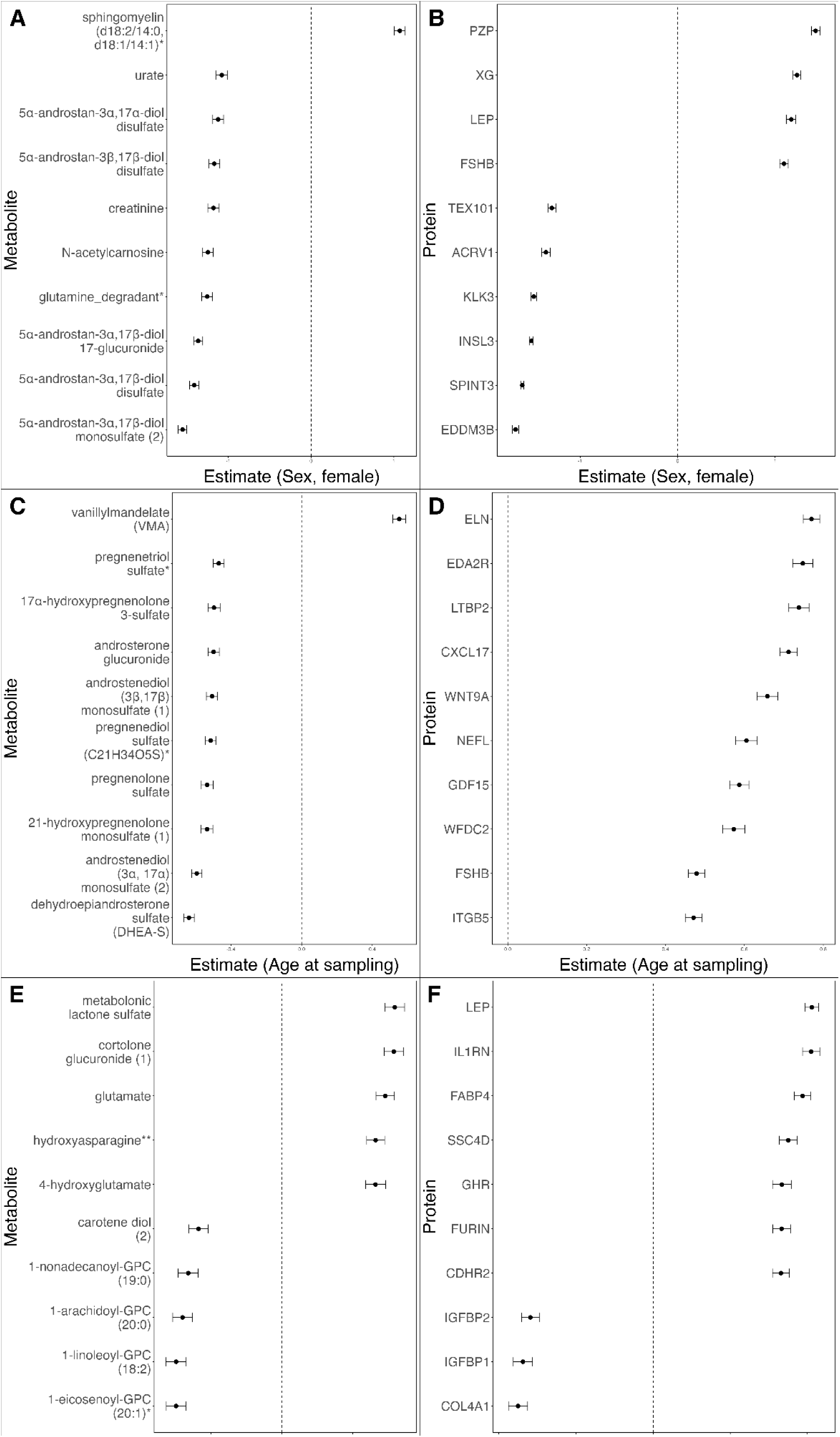
Forest plots of proteomics and metabolomics association analysis results with standard errors. Top ten associations are shown with the lowest FDR values for each covariate for metabolomics and proteomics. Metabolomics and proteomics values for female sex (A, B), age (C, D) and BMI (E, F).

**Supplementary Figure 3.**
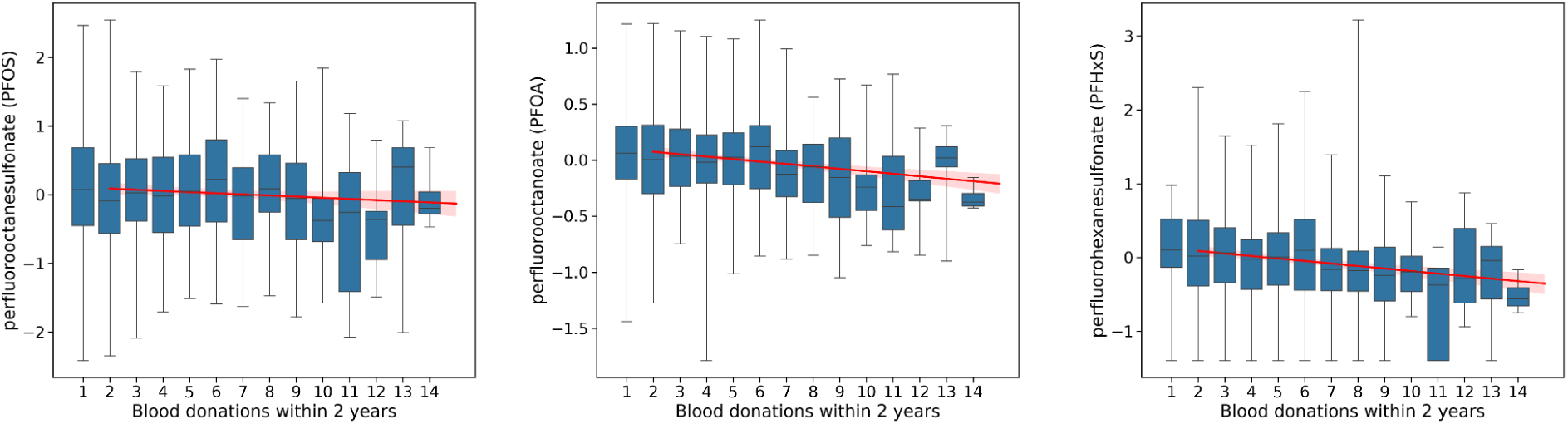
Levels of three PFAS compounds PFOS, PFOA and PFHxS show a decreasing trend with increasing numbers of blood donations.. Log transformed PFAS concentrations are plotted against blood donation frequency, with the fitted trend line shown in red.

## Supplementary Tables S1-S9

Association analysis results and other supplementary tables can be found in Supplementary Tables S1-S9.

## Supplementary Methods

### Genotyping of the samples

The biobank samples were genotyped as part of the FinnGen project with the FinnGen ThermoFisher Axiom custom array v1 or v2. Genotyping, quality control, and genome imputation protocols, R11, are described in detail in the FinnGen Gitbook (https://finngen.gitbook.io/documentation). In brief, genotype calling was performed with the AxiomGT1 algorithm. Prior to the imputation, genotyped samples were pre-phased with Eagle 2.3.5 with the default parameters, except the number of conditioning haplotypes was set to 20,000. Genotype imputation was performed using the population-specific imputation reference panel SISu v3 including 3,775 high coverage (25–30x) whole genome sequence data, with Beagle 4.1 (version 08Jun17.d8b). Genotypes were then returned to the Blood Service Biobank.

### Sample collection

The collection of samples from FinnGen-genotyped blood donors was done whenever they visited the Blood Service for their regular blood donation visit. No specific recalls were done. The sample collection was done in the Greater Helsinki area at three donation sites to ensure fast sample processing. All samples are being processed within 4 hours from the time of collection at the donation site. This is a critical time window for several of the downstream analyses of this study, especially the metabolomics and proteomics analyses. To increase the geographical coverage, ∼20% of the samples were collected at the blood donation sites of the cities of Oulu (North West Finland) and Kuopio (North East Finland).

Venous whole blood samples for plasma and serum aliquots were collected from the diversion pouch without anticoagulant immediately after blood donation. For plasma, two 4 ml EDTA tubes and one 4 ml serum tube were collected. Both plasma and serum tubes were centrifuged for 10 minutes at 2000 x g, and samples were subsequently aliquoted in 230 μL aliquots in FluidX 0.26 ml cryotubes with 2D barcodes (Cat. No. 68-0303-01). Aliquoting of plasma and plasma and serum was performed on a Hamilton Easy Blood Star liquid handling robot.

Venous whole blood samples were collected from the diversion pouch immediately after blood donation and used for harvesting the buffy coat (BC). Peripheral blood mononuclear cells (PBMCs) were isolated from BCs with Ficoll isogradient centrifugation. The BC preparation was diluted in 1:1 ratio with Dulbecco’s Phosphate Buffered Saline without Ca and Mg (DPBS, Gibco, Cat. NoA12856-01). The diluted blood was placed on top of 20 ml of Ficoll (GE Healthcare, Cat.No. 17-1440-03) in two 50 ml conical Falcon tubes. The tubes were centrifuged for 25 minutes at 800 x g (Eppendorf 5810R) at room temperature (RT) with no brake during the deceleration step. The upper plasma layer was removed by aspirating with a sterile glass Pasteur pipette. Then, the white layer containing the mononuclear cells was collected with a sterile individually-packed plastic Pasteur pipette. The cells were then transferred into a new 50 ml conical Falcon tube containing 40 ml DPBS. The tube was then filled to 50ml of DPBS and centrifuged at 300g for five minutes at RT. The supernatant was carefully removed by aspiration and the cell pellet was resuspended in 50 ml of DPBS and the cells were spun down as in the previous step. In total, the cells were washed three times with 50 ml of DPBS. After the washing steps, the cells were gently suspended in 3 ml of RPMI1640 cell culture medium supplemented with 5% of heat-inactivated human AB serum, 2mM L-glutamine and 1% of Pen-Strep. The cells were counted with an automated Bio-Rad TC20 cell counter or Burker chamber with trypan blue staining. To minimize variation across the samples the same serum batch was used for isolation and freezing of PBMC. The serum was aliquoted in 3-5 ml batches and kept frozen at −80 centigrades until used.

### Freezing and thawing protocols

For freezing, the density of the cells was adjusted to ∼2x10^7^ cells/ml. Then 5 ml of +4°C RPMI1640 cell culture medium with 5% of heat inactivated human AB serum, 2mM L-glutamine and 1% Pen-Strep and 20% of DMSO was added dropwise slowly while swaying the tube to gently mix the cell suspension (final solution: 10% DMSO in RPMI1640 medium supplemented with 5% inactivated human AB serum, 2mM L-glutamine and 1% Pen-Strep, herein freezing media). The cells suspended in the freezing medium were then mixed with gentle pipetting. Cell suspension was then aliquoted in the cryovials. One ml (1ml) of the cell suspension was transferred to a cryovial (FluidX 1.8ml screw cap tube with external threading). The tubes were closed and put immediately in a CoolCell LX (Corning) cell freezing container for temperature-controlled cooling of the cells. The CoolCell LX container was immediately transferred into a −80°C freezer overnight. The cryovials were transferred the next day into the automated gas phase liquid nitrogen freezer for long-term storing at -180°C.

The cryovials were quickly thawed in a heat-block pre-warmed at 37°C in an incubator. Once the cells began to thaw, warm RPMI1640 cell culture with 10% of heat inactivated human AB serum, 2mM L-glutamine and 1% of Pen-Strep (referred as thawing media from here on) was added on the top of icy cell suspension. Thawed cell suspension was repeatedly transferred in a 15 ml Falcon tube containing 10 ml of 37°C thawing media. The Falcon tubes were filled with 15 ml of thawing media and centrifuged for 10 minutes at 300 x g. The supernatant was carefully aspirated, and the pelleted cells were gently re-suspended in 5ml of regular culture media. Cells were centrifuged for 10 minutes at 300 x g and the supernatant was aspirated. Finally, the cells were re-suspended in 1 ml of regular cell culture medium and the cells were counted and viability of the cells was analyzed with an automated Bio-Rad TC20 cell counter or Burker chamber.

### Immune cell profiling

Thawed cells were stained with fluorochrome conjugated antibodies targeting cell surface antigens. For staining, cells were incubated at room temperature for 20 min, in dark, with a surface staining cocktail containing anti-CD4, anti-CD141, anti-CD123, anti- CD11c+, anti-HLA-DR, anti-CD127, anti- CD25 antibodies, and lineage cocktail 1 in a final volume of 150 µl. Subsequently, cells were washed once, resuspended in 0.5% BSA-PBS and kept on ice until FACS analysis. Analysis of 6 immune cell subpopulations (Treg-DC panel) was performed with BD Influx™ (BD Biosciences). Dead cells were excluded from analysis by SYTOX® Blue staining. The cell populations were defined as following: Teff (CD3+CD4+CD25intCD127+), Treg (CD3+CD4+CD25hiCD127-), mDC1, (HLADR+CD11c+CD141-), mDC2 (HLADR+CD11c+CD141+), (pDC: HLADR+CD123+CD4+) and monocytes (CD4lowHLA-DR+).

Gating strategy of the immune cell population is presented in Figure 3A. The list of antibodies and other reagents used in the FACS analysis is provided in the **Supplementary Table S9**.

### Immune cell activation and cytokine response

Frozen PBMCs were quickly thawed and resuspended in cell culture medium. PBMCs were stimulated for 48 h with plate-bound anti-CD3 (BD Pharmingen) and soluble anti-CD28 (1µg/ml) (BD Pharmingen). For T-cell stimulation, a 96-well cell culture plate (Nunclon) was pre-coated with anti-CD3 by incubating the cell culture plate with anti-CD3 at a concentration of 5µg/ml in PBS (50 µl in well) for 4 h at 37 °C in the presence of 5 % CO2. In total, six replicates of 2 x 10^5^ PBMCs were stimulated. The final density of the cells in the stimulation was 1x10^6^ / 1ml. After the stimulation, the cell culture supernatant was collected for cytokine analysis.

The serum concentrations of cytokines, chemokines and growth factors were analyzed using the 38-plexed Milliplex MAP Kit (cat.no. HCYTMAG-60K-PX38) according to the manufacturer’s recommendations (Merck-Millipore Corp., Billerica, MA, USA). Analyses were performed with single reactions. Quantification of the markers was performed with the Bio-plex 200 Luminex instrument and Bio-Plex Manager software (Bio-Rad, Sweden). Concentration of each marker was determined from an 8-point standard curve using five-parameter logistic regression. Minimum detectable concentration (MinDC) was determined for each marker separately using the lowest concentration on the standard curves linear phase (MinDC=c(low)+2SD). The samples below MinDC were given a value of 50% of MinDC. The following analytes were included in the 38-plex kit: sCD40L, EGF, Eotaxin, FGF2, FLT3L, Fractalkine, GCSF, GMCSF, GRO, IFNalpha, IFNgamma, IL1alpha, IL1beta, IL1RA, IL2, IL3, IL4, IL5, IL6, IL7, IL8, IL9, IL10, IL12p40, IL12p70, IL13, IL15, IL17A, IP10, MCP1, MCP3, CCL22, MIP1alpha, MIP1beta, TGFalpha, TNFalpha, TNFbeta, VEGF.

### Metabolomics

The samples were frozen within 4 hours after bleeding and stored at −80°C. Sample preparation was automated, including protein precipitation using methanol, followed by centrifugation. Extracts were divided into fractions for analysis under various chromatography and ionization methods—two reverse-phase UPLC-MS/MS with positive ion mode, one with negative ion mode, and one HILIC with negative ion mode. Quality assurance included spiked internal standards, pooled technical replicates (CMTRX), blanks, and solvent controls to assess and monitor variability across sample processing and instrumentation.

High-resolution mass spectrometry (UPLC-MS/MS) was conducted using a Thermo Q-Exactive Orbitrap system. Peak identification relied on a comprehensive library of over 3,300 authenticated standards and robust criteria, including retention index, accurate mass, and MS/MS spectral matching. Data processing involved stringent QC, curation to exclude artifacts, and quantification based on peak area. For multi-day analyses, normalization was corrected for batch effects using a block-correction method. Bioinformatics tools integrated LIMS tracking, automated data extraction, and visualization platforms to ensure high data fidelity for downstream statistical and biological interpretation.

### Proteomics

Data normalization was performed through a multi-step process designed to eliminate systematic biases and ensure consistency across samples and assay plates. The first step was Hybridization Control Normalization, which corrected for systematic effects during hybridization by using a set of control sequences added to each sample. Each sample’s data was scaled using a factor derived from the median ratio between control RFU values and a reference within the same plate. This was followed by Intraplate Median Signal Normalization, applied only to calibrator samples. Each SOMAmer reagent, grouped by dilution, was adjusted using a scale factor based on the median RFU of the reagents within that dilution, helping correct within-plate biases such as pipetting variation and reagent concentration differences. Next, Plate Scaling and Calibration addressed between-plate variability by comparing calibrator samples across plates to global reference standards established for each SOMAmer. A scale factor for each SOMAmer was calculated by dividing the global reference RFU by the local calibrator RFU on each plate, and this factor was applied across the entire plate. Finally, Median Signal Normalization to a Reference was performed on all individual, QC, and buffer samples by comparing each to a global reference dataset from healthy individuals. This step applied separate scale factors for each dilution group per sample, ensuring consistency across the full dataset. Together, these normalization steps reduced assay variability and standardized the data for robust downstream analysis.

